# Two components of the early ASH1 mRNA transport machinery undergo PUN motif-dependent liquid-liquid-phase separation

**DOI:** 10.64898/2026.01.20.700544

**Authors:** Wieland Mayer, Thomas Monecke, Juliane Bethmann, Henning Urlaub, Dierk Niessing

**Affiliations:** Institute of Pharmaceutical Biotechnology, Ulm University, Ulm, Germany; Max Planck Institute for Multidisciplinary Sciences, Bioanalytical Mass Spectrometry Group, Göttingen, Germany; University Medical Center Göttingen, Institute of Clinical Chemistry, Bioanalytics Group, Göttingen, Germany; Molecular Targets and Therapeutics Center, Institute of Structural Biology, Helmholtz Zentrum München - German Research Center for Environmental Health, Neuherberg, Germany

## Abstract

In eukaryotes, mRNA localization is a widespread mechanism of spatial gene regulation. Amongst the best-studied examples is the directional transport of *ASH1* mRNA during mitosis of the budding yeast *Saccharomyces cerevisiae*. *ASH1* mRNA is co-transcriptionally bound by the two RNA-binding proteins She2p and Loc1p and subsequently exported. In the cytoplasm, the adapter She3p binds to She2p forming an active transport complex with the myosin motor Myo4p to mediate actin-dependent transport into the daughter cell. Loc1p stays in the nucleus where it has a second function in ribosome biogenesis. In this study, we map the interaction surface of Loc1p on She2p and observe that it does not interfere with the RNA-binding interface of She2p. Analogous to a previous report on Loc1p and RNA, we could show that also Loc1p and She2p undergo liquid-liquid phase separation (LLPS). This event is caused by electrostatic interactions and can be regulated by the phosphorylation-driven alteration of She2p’s oligomeric state. Furthermore, we observed that LLPS formation only requires the PUN motifs of Loc1p and that a ternary Loc1p-RNA-She2p complex also undergoes LLPS. In summary, our findings indicate that Loc1p co-transcriptionally recruits the nuclear *ASH1* mRNA-She2p complex to LLPS, while the cytoplasmic Loc1p-lacking She2p-She3p-RNA transport complex does not form such LLPS.

## Introduction

Asymmetric mRNA localization is a process observed across eukaryotes (Das et al. 2021). It provides an opportunity to regulate, retain and transport transcripts in a spatial and temporal manner. To date, for only a limited number of transcripts cis-acting elements have been identified that are bound by a dedicated motor complex (Chekulaeva 2024). Even fewer examples are known where all protein factors required for the transport of a particular mRNA have been identified. The localization of the *ASH1* mRNA and about 30 other transcripts during mitosis of *Saccharomyces cerevisiae* presents an exceptionally well understood example of such a localization and the associated mRNP (Niessing et al. 2018). All essential factors have been identified, their interaction and stoichiometries been quantified, and the active transport of mRNA *in vitro* reconstituted from recombinant factors (Bobola et al. 1996; Sil and Herskowitz 1996; Gonzalez et al. 1999; Chartrand et al. 2002; Bohl et al. 2000; Heym et al. 2013; Müller et al. 2011; Jansen et al. 1996; Long et al. 2000; Takizawa and Vale 2000; Sladewski et al. 2013).

The transport process begins already co-transcriptionally with the dedicated RNA-binding protein (RBP) She2p recognizing cis-acting stem-loop structures of the emerging *ASH1* transcript (Shen et al. 2010). These cis-acting mRNA elements are termed localization or ZIP code elements and mediate the cytoplasmic attachment of transcripts to motor-protein-containing particles that move along actin filaments (Gonzalez et al. 1999; Chartrand et al. 1999, 2002). In addition to She2p, the nuclear factor Loc1p binds co-transcriptionally to the *ASH1* mRNA (Long et al. 2001; Shen et al. 2009) and is joined by the RNA-binding proteins Puf6p (Deng et al. 2008; Gu et al. 2004) and Khd1p (Irie et al. 2002a; Hasegawa et al. 2008; Paquin et al. 2007).

After a nucle(o)lar passage, the complex is remodeled at the nuclear pore (Du et al. 2007; Niedner et al. 2013). The cytoplasmic She3p adaptor protein, which is constitutively bound to Myo4p, displaces Loc1p from the complex (Bohl et al. 2000; Long et al. 2000; Niedner et al. 2013). The mature cytoplasmic complex consisting of the core factors She2p, She3p and the myosin motor Myo4p as well as its cargo mRNA actively moves along actin filaments to the bud tip of the emerging daughter cell. There, the mRNA is translated into Ash1p, which contributes to the prevention of mating type switching in the daughter cell (Cosma 2004).

Genomic deletion of the contributing factors indicated that She2p, She3p, and Myo4p are essential for *ASH1* mRNA transport (Jansen et al. 1996), whereas inactivation of Loc1p, Puf6p, and Khd1p has a moderate effect on this process (Hasegawa et al. 2008; Irie et al. 2002b; Paquin et al. 2007; Deng et al. 2008; Urbinati et al. 2006; Gu et al. 2004; Long et al. 2001). This *in vivo* observation is well reflected by *in vitro*-reconstitution experiments, showing that the essential functional transport complex consists of She2p, She3p, Myo4p and cargo RNA (Sladewski et al. 2013; Heym et al. 2013).

Since in *in vitro* binding assays She3p can displace Loc1p from its co-complex with She2p (Niedner et al. 2013) and because a sequence analysis revealed the presence of the (L)PGV(K) hook-motif of the cytoplasmic She3p also in Loc1p (**Figure 1A**), it seems likely that She3p and Loc1p use the same binding region in She2p for their complex formation.

**Figure 1:**
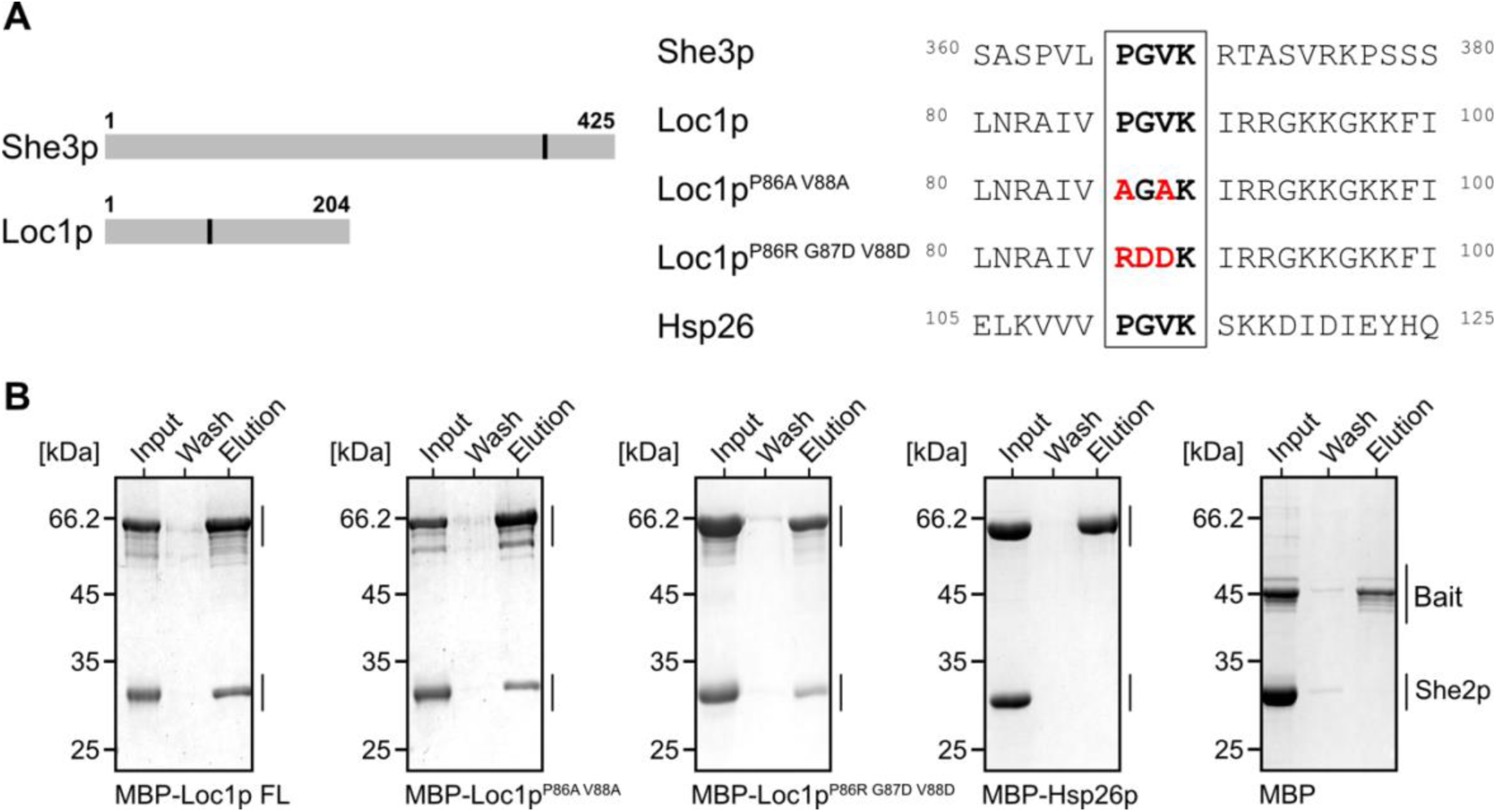
She2p interaction is not dependent on the PGVK-hook motif in Loc1p. **(A)** Position of the PGVK-motif in She3p and Loc1p and local sequence alignment of proteins used in this study. The PGVK (boxed) motif of She3p was shown to be required for its interaction with She2p and is also found in Hsp26p and Loc1p. Red letters indicate introduced mutations to probe the requirement of this motif in Loc1p for She2p binding. **(B)** Pull-down experiments using MBP-Loc1p, MBP-Hsp26p and MBP alone (control) as bait on amylose resin, showing that mutation of the PGVK motif does not impair the interaction of Loc1p and She2p. Vice versa, the presence of a PGVK motif is also insufficient for interaction in the case of Hsp26p. Experiments were performed in triplicates (n = 3).

In addition to the above-described function in mRNA transport, Loc1p has been implicated in ribosome biogenesis. For instance, Loc1p was co-immunoprecipitated with 66S and 90S particles (Jung et al. 2019; Liang et al. 2019; Urbinati et al. 2006) and *loc1* deletion strains display a slow-growth phenotype and reduced 60S levels, suggesting a role of Loc1p in ribosome maturation (Niedner-Boblenz et al. 2024). A detailed assessment of the rRNA processing steps revealed reduced levels of 20S and 18S pre-rRNA and an accumulation of 35S pre-rRNA (Urbinati et al. 2006). Moreover, cryo-EM structures of *in vivo* isolated pre-60S particles showed directly that Loc1p is associated with late-nucleolar intermediates of ribosomal biogenesis (Cruz et al. 2022). While these findings described the consequences of impaired Loc1p function on nucleolar processes, they fell short of providing insights into the molecular functions of Loc1p.

Only very recently, it was demonstrated that Loc1p mediates RNA annealing as well as liquid-liquid phase separation (LLPS) (Niedner-Boblenz et al. 2024), features that are both required for rRNA processing in the nucleolus (Dignon et al. 2020). The study further identified several linear sequence elements in Loc1p, so-called <u>p</u>ositively charged <u>u</u>nfolded <u>n</u>ucleic acid-binding (PUN) motifs, that act cooperatively to achieve RNA annealing.

While RNA annealing and LLPS are both features of Loc1p likely required for ribosomal maturation in the nucleolus, it remains unclear if they also impact the second function of Loc1p, i.e. the assembly of nucle(o)lar complexes for subsequent cytoplasmic *ASH1* mRNA transport. In this study, we present insights into how Loc1p interacts with the non-canonical RBP She2p. We demonstrate that Loc1p, She2p, and nucleic acids can undergo LLPS and that PUN motifs in Loc1p are necessary and sufficient for this function. Hence, our results indicate that LLPS via multivalent interactions may be important for the assembly of transport pre-complexes in the nucleus but rather play no role in the cytoplasmic *ASH1* mRNA-transport machinery.

## Results

### The conserved PGVK motif of Loc1p does not mediate the Loc1p-She2p interaction

Since the She2p-interacting LPGV(K) motif of She3p is conserved in Loc1p (**Figure 1A**), we first assessed whether this motif is also responsible for the Loc1p-She2p interaction. We performed *in vitro* pull-down assays with recombinant MBP-tagged wild type Loc1p or variants bearing mutations in its PGVK motif, and She2p as prey. The Loc1p variants contained either conservative mutations in the PGVK motif (Loc1p^P86A V88A^) or more dramatic exchanges that altered side chain lengths and charges, leading to potential steric clashes (Loc1p^P86R G87D V88D^). In the past it was shown that the introduction of only one of the latter single point mutations in She3p (P365R or G366D or V367D) was sufficient to disrupt its interaction with She2p (Singh et al. 2015). In contrast, we did not observe any impairment of the She2p-Loc1p interaction even when all three mutations were combined in Loc1p^P86R G87D V88D^ (**Figure 1B**). We concluded that the PGVK motif in Loc1p is not required for its interaction with She2p.

### Multiple Loc1p regions mediate the interaction with She2p

Loc1p was recently shown to lack folded domains and tertiary structure (Niedner-Boblenz et al. 2024). In order to map the regions of Loc1p that interact with She2p, we used a tiling array consisting of 20 aa-long Loc1p peptides with a shift of 3 aa between each peptide and performed a binding experiment with recombinant She2p (**Supplementary Figure 1**). Quantification of normalized spot intensities allowed us to identify Loc1p peptides 13-20, 25-31, and 52-58 that bound to She2p, suggesting that Loc1p contains three main She2p-interacting regions (**Figure 2A-C**). Interestingly, these three regions partially overlap with previously identified stretches of Loc1p that bound to RNA and had been termed PUN motifs (Niedner-Boblenz et al. 2024).

**Figure 2:**
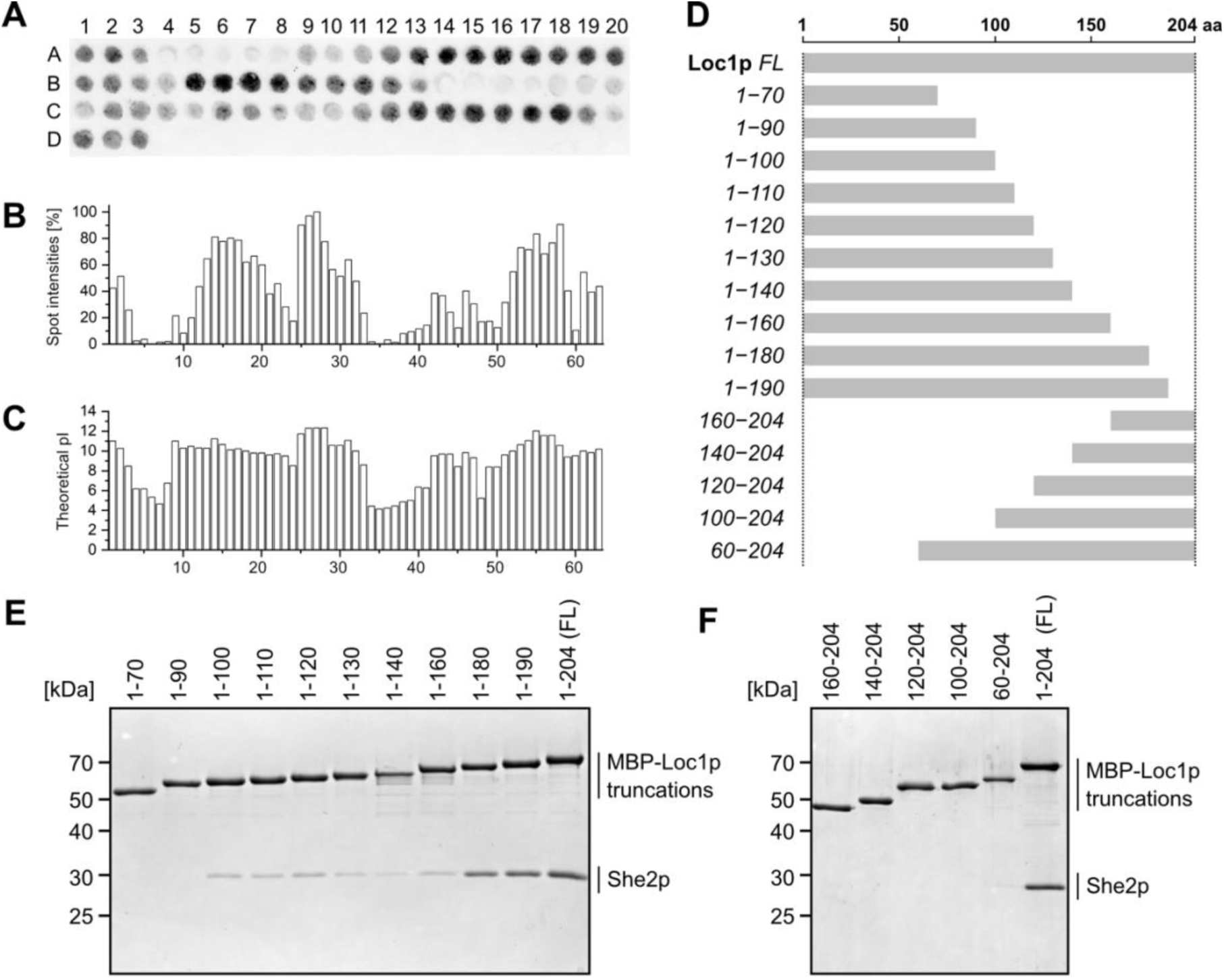
Multiple regions of Loc1p interact with She2p. **(A)** Peptide tiling array revealed binding of She2p to 20 aa Loc1p peptides, each shifted by 3 aa. Spots were incubated with recombinant She2p, which was detected by a primary anti-She2p (rat) antibody and a secondary anti-rat IRDye^®^ 680 RD (n = 1). **(B)** The spot intensities were normalized to 100% (strongest spot defined as 100%) and plotted against the respective spot number, revealing a pattern of multiple She2p-binding regions. **(C)** The spot number was plotted against the theoretical isoelectric point (pI) of the peptides, showing a clear correlation between binding intensities and positive charge. **(D)** Schematic representation of Loc1p constructs used in the pull-downs experiments (see below). **(E)** SDS-PAGE of the elution fractions of *in vitro* pull-down experiments, using C-terminal MBP-Loc1p truncations as bait and She2p as prey. **(F)** N-terminal MBP-Loc1p deletions that were tested as in (E) for binding to She2p, revealing a complete loss of binding upon truncation. Additional controls for all pull-down experiments, including input and last wash fractions, are shown in **Supplementary Figure 2.**

Since the peptide tiling array only gives information about the binding of individual Loc1p peptides to She2p, we also tested the interaction using truncated Loc1p fragments in *in vitro* pull-down experiments. While increasing C-terminal truncations of Loc1p resulted in a progressive loss of She2p co-elution, N-terminal truncation led to a complete loss of binding (**Figure 2D-F**). From these experiments, we conclude that large parts of Loc1p are required for She2p interaction. Combined with the observations from the tiling array, these insights suggest the requirement of multiple binding sites in Loc1p for efficient She2p interaction.

### Loc1p is a monomeric protein that forms LLPS with She2p

The stoichiometry of the mature cytoplasmic mRNA complex is well understood. A tetramer of She2p forms the core of the mRNP, which binds two mRNA localization elements on opposing sides of the molecule (Niessing et al. 2004; Müller et al. 2009; Edelmann et al. 2017), and four molecules of She3p, which interact as two dimers with Myo4p (Krementsova et al. 2011; Heym et al. 2013; Edelmann et al. 2015). To understand if the nuclear complex consisting of She2p and Loc1p adopts the same 1:1 stoichiometry as She2p and She3p, we performed multi-angle light scattering (MALS) measurements. Loc1p alone remained monomeric even at a concentration of 115 µM (**Supplementary Figure 3A, B**). As previously reported, She2p alone formed a tetramer in solution (Edelmann et al. 2017; Niessing et al. 2004) (**Supplementary Figure 3D**). The monomeric state of Loc1p was further verified using pull-down experiments with MBP-Loc1p as bait and His_6_-Loc1p as prey. No interaction could be detected by this orthologous approach, confirming the monomeric nature of the protein in solution (**Supplementary Figure 3B**). MALS measurements of She2p-Loc1p to determine the complex stoichiometry were hampered by the fact that the solution became cloudy immediately after mixing both proteins. Initially, we interpreted the observed effect as precipitation since the solution could be cleared by centrifugation and the supernatant was free of protein. However, considering the recently reported phase-separation of Loc1p with RNA *in vitro* (Niedner-Boblenz et al. 2024) we revisited the observed turbidity. Although in this earlier report, the presence of RNA was essential for LLPS formation, we wondered if also a protein like She2p could drive phase separation with Loc1p. Indeed, while no such droplets formed with the individual proteins at concentrations of 10 µM, mixing both proteins at the same concentrations resulted in round, floating entities (**Figure 3A**). This observation suggests that Loc1p and She2p together likely form LLPS.

**Figure 3:**
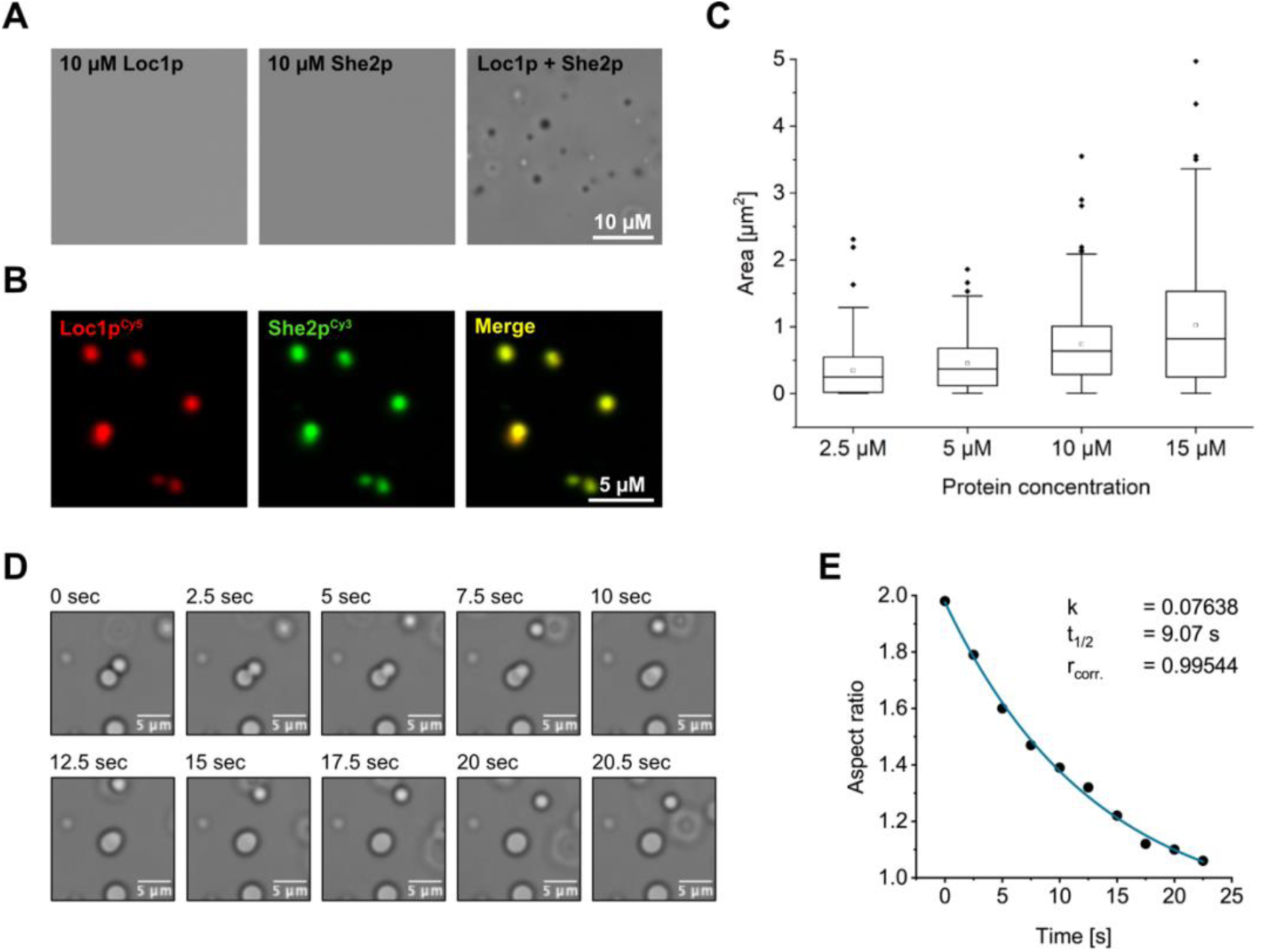
She2p and Loc1p form liquid-liquid phase-separated condensates. **(A)** Mixing recombinant His_6_-Loc1p and She2p *in vitro* resulted in the formation of LLPS entities. Proteins at equimolar concentrations of 10 µM were mixed in buffer with 150 mM NaCl and imaged under a light microscope after 5 min of incubation at room temperature. No LLPS-inducing agents like PEGs were used for LLPS experiments. **(B)** Imaging of fluorescently labeled proteins showed that observed droplets consist of both proteins. **(C)** Quantification of droplets from light-microscopic images revealed that droplet sizes correlate with the corresponding protein concentration. Proteins were mixed at various concentrations in a buffer containing 150 mM NaCl and imaged after 5 minutes of incubation at room temperature. **(D)** Time-resolved images of two droplets fusing over a time of 20.5 sec. **(E)** The aspect ratio of perpendicular axes from a representative fusion event was plotted against time and showed a decrease from 2.0 (elliptic) to 1.0 (round), illustrating the liquid character of the fusing phases. The exponential decay function revealed a rate constant (k) of 0.07638 s^-1^ and a half-life time of 9.07 sec. (rcorr of the fit was calculated as 0.99544).

To directly demonstrate the presence of both proteins in LLPS entities, we mixed fluorescently labeled Loc1p^Cy5^ and She2p^Cy3^ and imaged the resulting droplets by fluorescence microscopy. Both proteins colocalize to the same phase-separated droplets as observed by their overlapping fluorescent signals (**Figure 3B**). In addition, we observed that the sizes of the entities positively correlate with the protein concentration (Figure 3C) (**Figure 3C**). To further assess the liquid-like nature of these LLPS, we characterized them in a coalescence assay where the condensates clearly fused and relaxed into spherical shapes after fusion (**Figure 3D,E** and **Supplementary Movies 1** and **2**).

To better understand how Loc1p interacts with She2p in the droplets, we applied chemical carbodiimide cross-linking followed by enzymatic digestion and mass-spectrometry (XL-MS) **(Supplementary Figure 4** and **Supplementary Tables 1-3)**. The amino acids of She2p that cross-linked to Loc1p were plotted onto the crystal structure of She2p (PDB-ID 5M0I) and are either located laterally at the She2p tetramer or at its top and bottom (**Figure 4A**).

**Figure 4:**
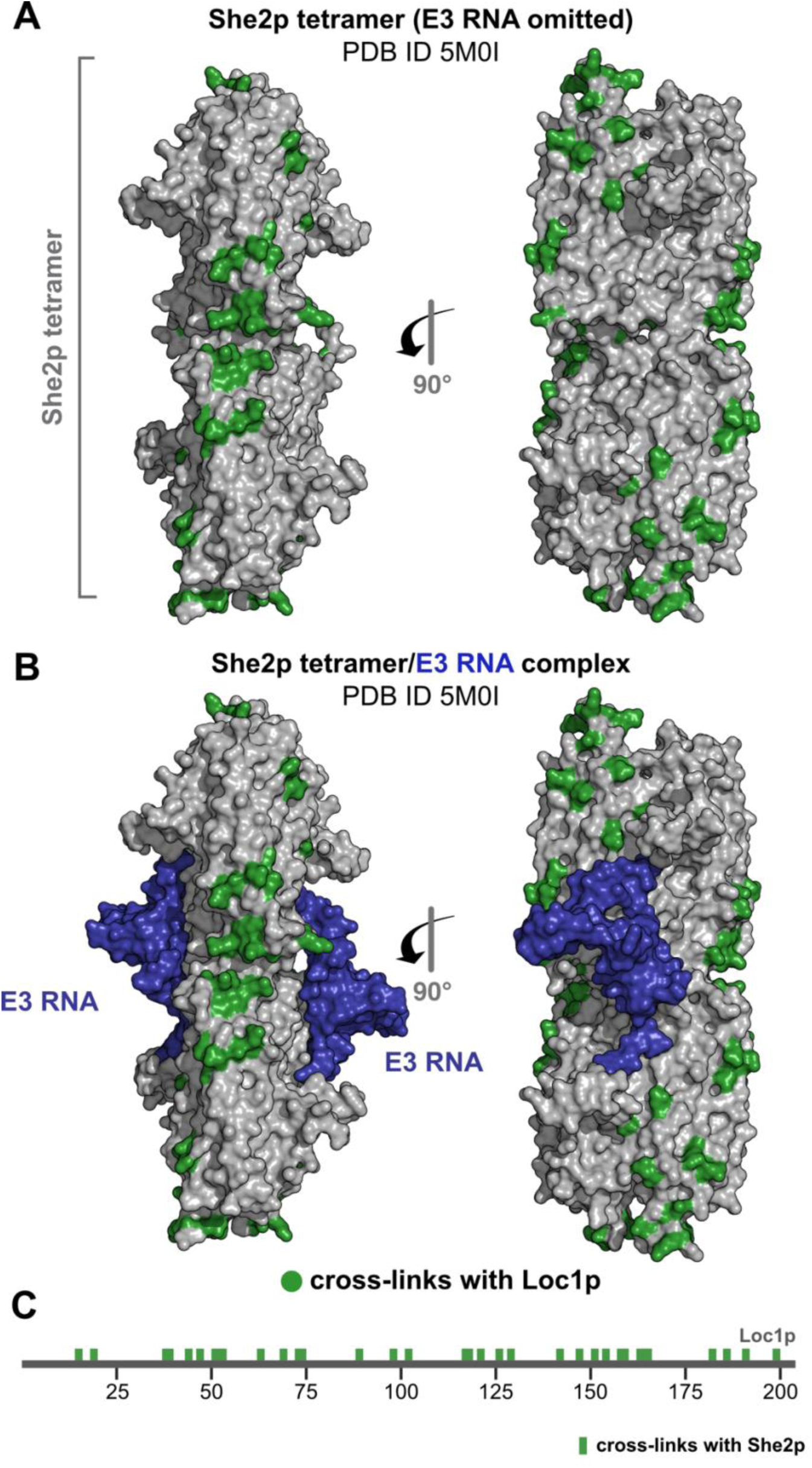
Cross-linking experiments with the Loc1p-She2p co-complex under LLPS-forming conditions. **(A)** Amino acids of She2p cross-linking to Loc1p (green) were plotted onto the crystal structure of She2p (PDB ID 5M0I). **(B)** The same crystal structure as in (A), with the bound E3 ZIP code RNA shown in blue and Loc1p cross-links indicated in green. Note that the RNA-binding surface of She2p does not overlap with Loc1p-cross linking sites. **(C)** Plot of the peptides in the primary sequence of Loc1p that cross-linked to She2p. Cross-links are distributed in clusters over the whole Loc1p sequence.

Previous results showed that She2p forms a ternary complex with RNA ZIP codes and Loc1p (Niedner et al. 2013), suggesting that the binding regions for both interactors do not overlap. Indeed, when visualizing the cross-linked Loc1p peptides together with the E3 RNA (PDB-ID 5M0I) (Edelmann et al. 2017), no overlaps of the respective interaction surfaces of the RNA and Loc1p were observed (**Figure 4B**). This observation does not only validate the specificity of the cross-linking results but also provides spatial information about the ternary complex of She2p, ZIP-code RNA and Loc1p. Since the protein-protein XL-MS experiment was performed under LLPS-forming conditions and in the absence of a ZIP code RNA, the non-overlapping Loc1p- and RNA-interaction surfaces on She2p show that the She2p-Loc1p interaction sites are specific, also in LLPS.

Since Loc1p consists of largely unstructured regions (Niedner-Boblenz et al. 2024), its interaction with She2p cannot be mediated by folded domains. When plotting the cross-linked amino acids of Loc1p onto its primary sequence, we observed a clustered distribution of the cross-links over the protein sequence (**Figure 4C**). This is in line with the observations made in the peptide tiling array (**Figure 2A-C**) that also show several regions or clusters interacting with She2p. Moreover, it is also consistent with the pull-down assays, where neither the N-terminal nor the C-terminal half alone stably interacted with She2p (**Figure 2E,F**).

### LLPS is mediated by electrostatic interactions and requires She2p oligomerization as well as PUN motifs in Loc1p

To understand the nature of the LLPS-driving interactions, we probed the She2p-Loc1p droplets using 1,6-hexanediol. This aliphatic alcohol has been shown to disrupt LLPS *in vitro* and biomolecular condensates *in vivo* if they are mediated by weak hydrophobic interactions. Formation of She2p- and Loc1p-containing droplets at and above physiological concentrations (1.25 µM and 10 µM, respectively) were resistant to 5% and 10% of 1,6-hexanediol (**Figure 5A**). Hence, formation of the observed Loc1p-She2p condensates is likely not driven by hydrophobic interactions. Considering that She2p-interacting regions of Loc1p are particularly positively charged, (**Figure 2A-C**), it seems rather likely that LLPS formation is driven by charged, hydrophilic interaction.

**Figure 5:**
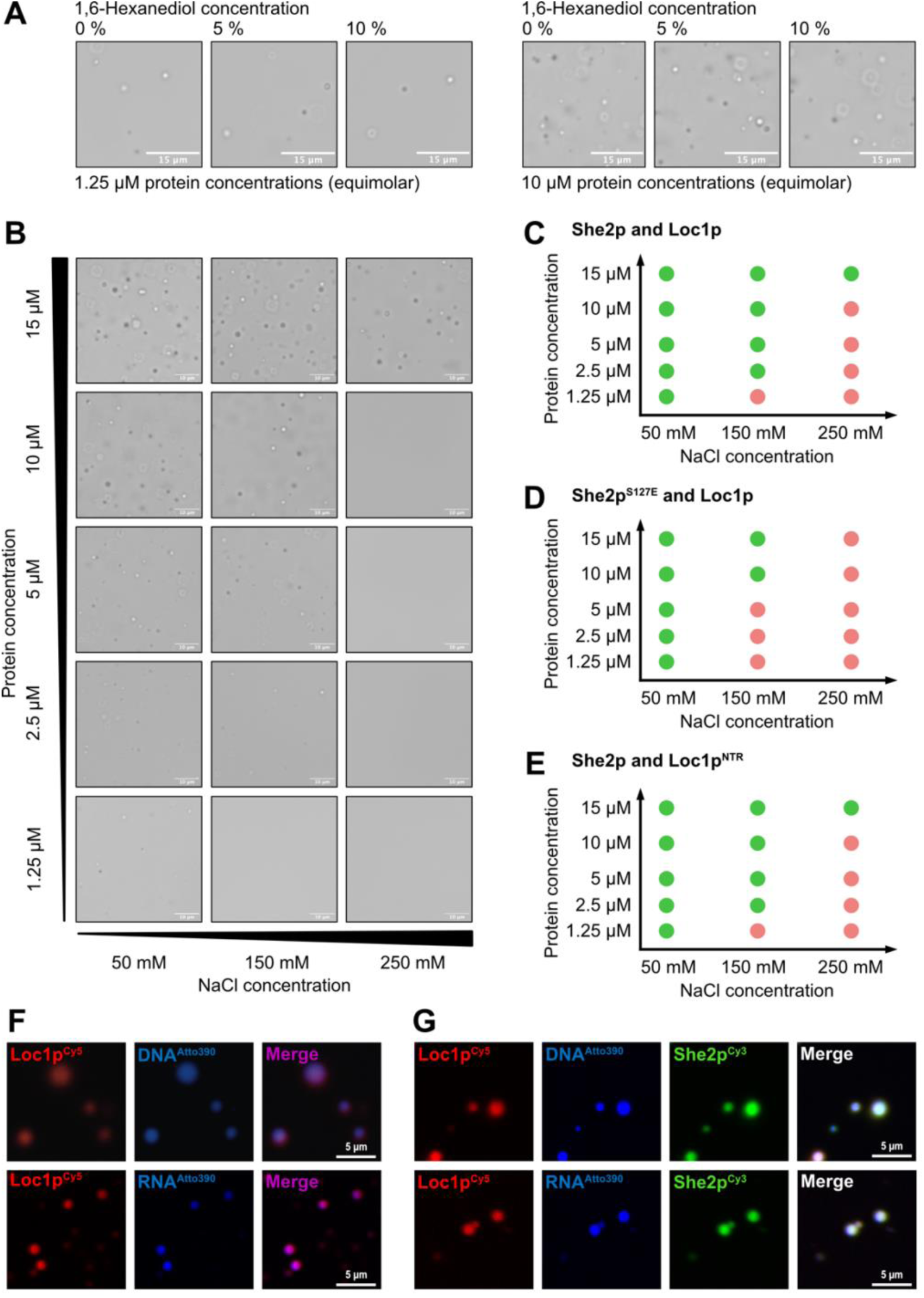
Characterization of LLPS properties of She2p-Loc1p and She2p-Loc1p-nucleic acid complexes. **(A)** Weak hydrophobic interactions are unlikely to cause LLPS. The addition of 5% or 10% 1,6-Hexanediol does not impair the formation of liquid condensates, ruling out weak hydrophobic interactions as the driving force for LLPS. Buffer and 1,6-Hexanediol were mixed before the individual proteins were added. Phases were imaged after 5 min incubation at room temperature with either 1.25 µM (left) or 10 µM (right) concentration of each protein in the co-complex. **(B)** Representative light-microscopy images of phase-separated She2p-Loc1p droplets at different protein and NaCl concentrations. **(C)** Phase diagram of She2p (wild-type, tetramer) and Loc1p, derived from images shown in (B) with n=3. The formation of phase-separated droplets is dependent on protein concentration and ionic strength (NaCl concentration). **(D)** Dimeric She2p (S127E) shows an altered LLPS phase diagram, with no droplets visible at 250 mM NaCl. **(E)** PUN motifs of Loc1p are sufficient to drive LLPS with She2p. A Loc1p construct consisting of 10x the 20 N-terminal amino acids containing the first PUN-motif of Loc1p (<u>N</u>-terminal <u>r</u>epeat, NTR) shows a Loc1p wild type-like phase diagram (compare with C). All experiments were performed in duplicates. **(F)** Cy5-labeled Loc1p alone forms LLPS with either Atto390-labeled DNA or the E3 RNA ZIP code. **(G)** Cy5-labeled Loc1p and Cy3-labeled She2p form LLPS with Atto390-labeled nucleic acids. First, both proteins were mixed at equal concentrations and either a labeled ZIP-code RNA or a labeled DNA-oligonucleotide was added afterwards. In both cases, all three components are present in the phase-separated droplets.

To challenge this hypothesis, we tested whether the increase of ionic strength influences phase-separation of equimolar Loc1p/She2p mixtures. At 1.25 µM protein concentration, we observed droplet formation at 50 mM NaCl but not at higher salt concentrations (**Figure 5B, C**). At higher protein concentrations (2.5, 5 and 10 µM), LLPS formed at 150 mM but not at 250 mM NaCl. Only at the 15 µM, the highest protein concentration tested, LLPS formation was observed at 250 mM NaCl, further supporting the hypothesis that She2p-Loc1p LLPS formation is mediated by electrostatic interactions.

Since multiple parameters could impact LLPS, including oligomeric states of the involved proteins, we assessed whether She2p needs to be tetrameric for LLPS formation with Loc1p. For this, we utilized a mutant version of She2p (S127E), which bears a point mutation in the tetramerization interface and results in dimeric She2p (**Supplementary Figure 3C, D**). When She2p (S127E) was tested for LLPS formation with Loc1p, we observed a reduced propensity to form LLPS (**Figure 5D**). This difference was most clearly observed at medium and higher ionic strengths. It indicates that the oligomeric state of She2p has an impact on its ability to contribute to LLPS formation. In fact, the observed lower tendency for LLPS formation with the dimeric She2p^S127E^ is in line with the concept of multivalent interactions required for efficient LLPS. The possibility for electrostatically driven multivalent interactions is lower for a She2p dimer compared to a tetramer interacting with Loc1p. Interestingly, such changes of the oligomeric state of She2p have been suggested to be caused by phosphorylation (Farajzadeh et al. 2023). Such a regulation would be suitable to change the propensity of She2p to form biomolecular condensates with Loc1p in the nucleus.

In the peptide tiling array (**Figure 2A-C**), many regions of Loc1p bound to She2p. These regions overlap with the previously identified RNA-binding PUN motifs in Loc1p (Niedner-Boblenz et al. 2024). Since these PUN motifs were recently shown to mediate LLPS formation of Loc1p with nucleic acids, we investigated the role of PUN motifs in Loc1p for LLPS formation with She2p. To uncouple the overall composition and functions of the full-length Loc1p from the multivalent interaction principle of PUN motifs, we resorted to a previously described synthetic protein consisting of a ten-times concatenation of the N-terminal 20 amino acids, containing the first PUN motif of Loc1p. By utilizing this Loc1p^NTR^ (NTR = <u>N</u>-terminal <u>r</u>epeat) protein with She2p for LLPS formation, we observed a phase diagram almost indistinguishable from the same experiment with wildtype Loc1p (**Figure 5E**). This observation suggests that clustered PUN motifs are sufficient to drive the LLPS with She2p. Taken together, we could show that the combination of PUN-driven electrostatic interactions of Loc1p with tetrameric She2p are prerequisites for the formation of LLPS at physiological concentrations.

### Incorporation of nucleic acids into phase-separated droplets

The nucleolus is a phase-separated, nuclear biomolecular condensate formed around rDNA and rRNA (Schwarzacher and Wachtler 1993; Lafontaine et al. 2021). Our current understanding of *ASH1* mRNA localization assumes that prior to a nucleolar transition of the transcript, co-transcriptional binding of She2p and Loc1p to the nascent *ASH1* mRNA occurs (Shen et al. 2010). To understand if this ternary complex would also have the propensity to undergo LLPS, we tested whether Loc1p- and She2p-containing phase-separated droplets also incorporate nucleic acids. By fluorescently labeling the individual components, we observed that Cy5-labeled Loc1p (Loc1p^Cy5^) forms LLPS with short (33 nt) Atto390-labeled DNA oligonucleotides or a 28 nt (E3) mRNA ZIP code element (**Figure 5F**). In the next step, Loc1p^Cy5^ was pre-incubated with Cy3-labelled She2p (She2p^Cy3^) before either Atto390-labeled DNA oligonucleotides or mRNA ZIP code element was added. Fluorescence imaging revealed that both, RNA and DNA oligonucleotides were incorporated into the pre-formed She2p^Cy3^/Loc1p^Cy5^ liquid droplets (**Figure 5G**). In summary, these findings suggest that She2p, Loc1p, and ZIP code mRNA can form phase-separating ternary complexes and that RNA can be recruited to pre-existing She2p/Loc1p phases.

## Discussion

Previous studies already examined the interaction of She2p and Loc1p and showed that the nucleolar passage of the *ASH1* mRNP (Du et al. 2008; Shen et al. 2009) influences the transcripts’ cytoplasmic transport along actin filaments towards the bud tip of the daughter cell. Such nucleolar passage is not unique to *ASH1* mRNA as for instance also the mammalian RNA-binding proteins Staufen 1 and 2 were shown to accumulate in the nucleolus (Martel et al. 2006; Macchi et al. 2004). This step is considered to be important for the biogenesis of the ribonucleoprotein complex, for its nuclear export and further transport. For the transport of *ASH1* mRNA by the SHE-proteins in yeast, the cytoplasmatic part of the transport process has already been studied in great detail. For instance, crystallographic studies characterized the interaction of She2p and the cytoplasmic myosin adapter She3p, and determined an (L)PGVK-hook motif on She3p to be important for this interaction (Singh et al. 2015; Edelmann et al. 2017). Our initial hypothesis was that also the interaction of She2p and Loc1p could be mediated by a similar, conserved PGVK sequence present in Loc1p. We applied *in vitro* pull-down assays and did not observe a significant reduction of bound She2p to MBP-Loc1p when mutating the PGVK-motif in Loc1p. We cross-validated our finding that a PGVK motif by itself does not mediate She2p binding with the unrelated yeast heat shock protein Hsp26p. A peptide tiling array of Loc1p and *in vitro* pull-down assays using several Loc1p-deletion mutants revealed instead that multiple regions of Loc1p interact with She2p.

Since Loc1p contains predicted nuclear and nucleolar localization signals and because She2p lacks such sequences, a piggy-back import of She2p into the nucleolus by Loc1p was assumed (Niedner et al. 2013). A major result of the present study is that She2p and Loc1p undergo liquid-liquid phase separation (LLPS). When estimating the physiological concentration of Loc1p in the nucleus to be in the range of 2.5-3.0 µM (for calculation, see Material & Methods), it indicates that the protein concentration of Loc1p used in our *in vitro* experiments (see **Figure 5**) was sub-physiological. Together these considerations suggest that the LLPS-forming property may contribute to the correct localization of the complex to the nucleolus, which is itself a phase-separated entity. Although we observed that a protein construct with ten concatenated N-terminal PUN motifs (Loc1p (NTR)) was able to drive LLPS with She2p, it remains to be shown if PUN motifs alone would also be able to mediate nucleolar sub-localization of the complex.

To better understand the interaction of Loc1p and She2p under phase-separated conditions, we performed in-phase XL-MS experiments. The mapped binding regions of Loc1p on the surface of the She2p tetramer do not overlap with its RNA-binding, indicating specific interactions of both proteins also under LLPS conditions. Thus, the XL-MS experiments provide important information on the spatial arrangement of the ternary She2p-Loc1p-RNA complex. When mapping the She2p-crosslinked amino acids on Loc1p, we uncovered that most cross-linked sites are enriched in the N-terminal half of the protein. This finding is consistent with our observation that N-terminal truncations of Loc1p completely abolished the interaction with She2p.

LLPS formation can rely on different principles of multivalent molecular interactions. We observed that the interaction of She2p with Loc1p and RNA in phase-separated droplets is based on electrostatic interactions and can be modulated by the ionic strength of the environment. With this feature, the droplets follow the same principles as other well-studied proteins undergoing LLPS such as p53 (Kamagata et al. 2020), and the tau protein (Boyko et al. 2019). Additionally, the oligomeric state of She2p influences the propensity to form LLPS droplets with Loc1p at physiologic protein concentrations (**Supplementary Figure 3C, D**). This observation indicates a potential She2p oligomerization-dependent regulatory mechanism of LLPS formation in yeast.

Indeed, Farajzadeh and colleagues recently showed that She2p is post-translationally modified by phosphorylation, impacting the asymmetric distribution of the *ASH1* mRNA (Farajzadeh et al. 2023). A phospho-mimetic mutation of the corresponding amino acid (T109D) disrupted the tetrameric state of She2p and impaired its interaction with the myosin adaptor She3p and the importin-α Srp1p. Of note, it has been previously shown that She2p is stable as a tetramer only at µM concentrations (Müller et al. 2009). In light of the regulatory role assigned to the oligomeric state of She2p, it seems reasonable to assume that the moderate stability of She2p might be a prerequisite for the phosphorylation-induced changes in oligomeric state of She2p. Together with our results, these findings suggest that changes in the oligomeric state of She2p via post-translational modifications do not only alter its interactions with transport factors but also modulates its propensity to form LLPS with Loc1p in the nucle(ol)us.

Understanding the features of LLPS formation by Loc1p, She2p and RNA expands our view on Loc1p-related events in the nucle(ol)us. This feature provides a rationale for the nucleolar enrichment of Loc1p for ribosome biogenesis. It is also suited to explain why She2p and ZIP code RNA enters the nucleolus with Loc1p (**Figure 6**), thereby indicating that Loc1p uses its biophysical properties of LLPS formation similarly for its two independent functions. It has been shown that co-transcriptional binding of She2p to nascent mRNAs is mediated by the RNA polymerase II elongation factor Spt4/5 (Shen et al. 2010). Moreover, a recent study demonstrated that phase-separation contributes to the formation of RNA polymerase II clusters in the nucleolus (Rippe and Papantonis 2022). Hence, the ability of Loc1p to induce LLPS with She2p and RNA might contribute to a greater organizational principle involving transcription and the nucleolus.

**Figure 6:**
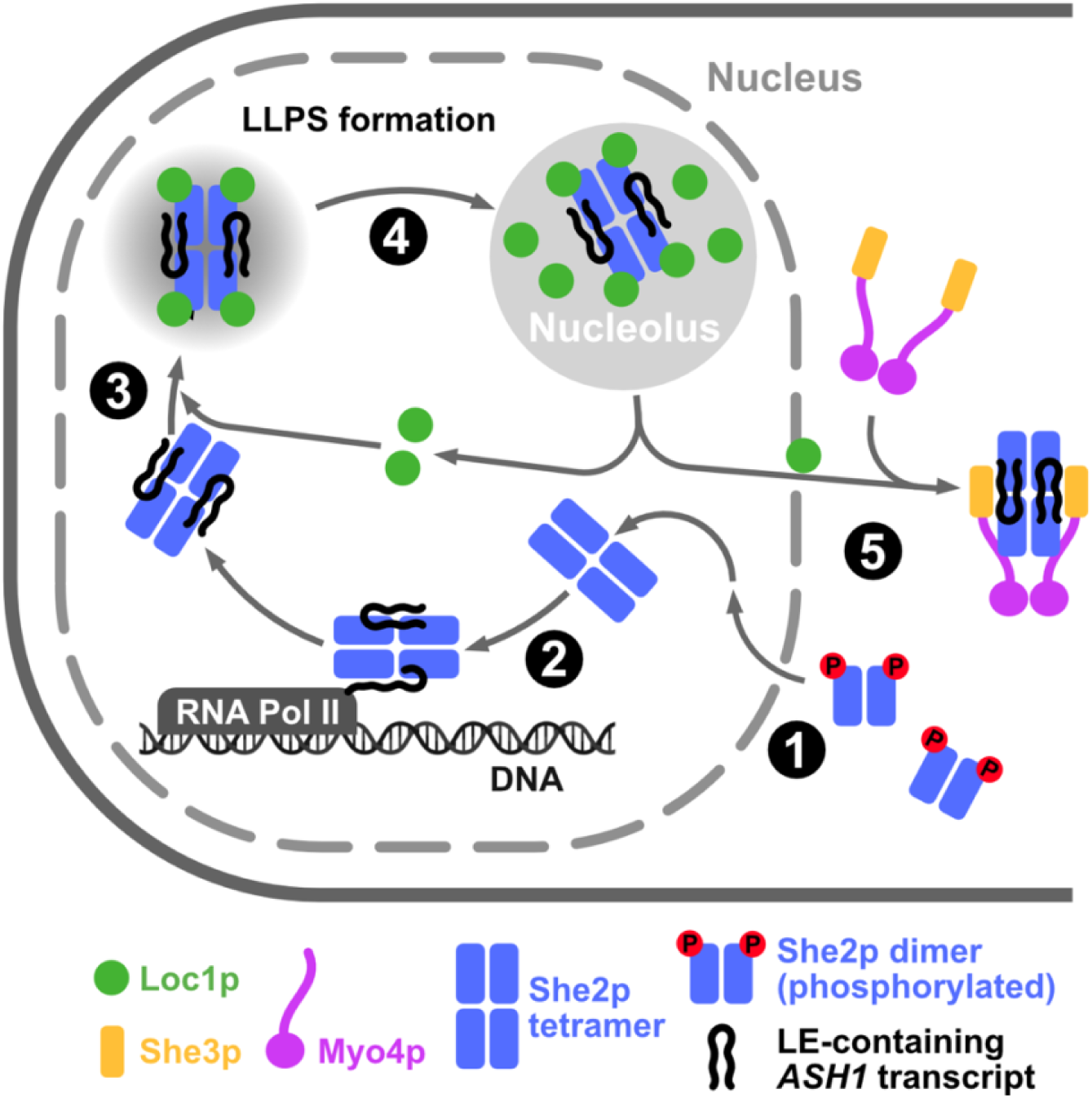
Model of LLPS-assisted nuclear mRNP-maturation and assembly. (1) She2p in a lower oligomeric state (most likely caused by phosphorylation) interacts with Srp1p (not shown) and is imported into the nucleus. To efficiently bind ZIP code-containing transcripts, She2p would need to get dephosphorylated and reassemble into a tetramer. **(2)** She2p tetramer is co-transcriptionally recruited to the emerging transcript and binds ZIP code-containing transcripts. **(3)** She2p may undergo LLPS together with Loc1p and RNA in the nucleus. The tetrameric state of She2p, which is a pre-requisite of efficient ZIP code-RNA binding, also promotes LLPS formation with Loc1p. **(4)** Phase separated nuclear droplets localize correctly assembled nuclear pre-mRNP to the nucleolus. **(5)** During the re-location of the mRNP to the cytoplasm, Loc1p is released from the complex and replaced by She3p. For this, a handover between Loc1p and She3p at the nuclear pore is the most likely scenario.

## Materials and methods

### Protein expression and purifications

MBP-Loc1p and MBP-Loc1p truncations were expressed in *E. coli* BL21 (DE3) from a pET-M43 vector and in 2YT media containing 0.2% glucose. Cultures were grown to an OD_600_ of 0.8, and expression was induced with 0.25 mM IPTG. Induced cells were grown at 18 °C for 16 hours before being harvested and flash-frozen in liquid nitrogen. Cells were resuspended in lysis buffer (20 mM HEPES/NaOH pH 7.5, 500 mM KCl, 0.5 mM EDTA) together with c0mplete protease inhibitor (Roche) and lysed using a microfluidizer and cleared by centrifugation. Supernatant was applied to an amylose resin column and extensively washed with lysis buffer, high-salt buffer (20 mM HEPES/NaOH pH 7.5, 1000 mM KCl, 0.5 mM EDTA) and low-salt buffer (20 mM HEPES/NaOH pH 7.5, 150 mM KCl, 0.5 mM EDTA) and eluted with low-salt buffer containing 20 mM D-maltose. Eluted protein from affinity chromatography was bound to a heparin column (HiTrap™ Heparin HP, Cytiva) and eluted using a linear KCl-gradient (20 mM HEPES/NaOH pH 7.5, 1000 mM KCl, 0.5 mM EDTA). Final polishing was done using a HiLoad™ 16/600 Superdex™ 75 pg gel filtration column (Cytiva) in 20 mM HEPES/NaOH pH 7.5, 150 mM NaCl and 0.5 mM EDTA.

MBP-Hsp26p was expressed in *E. coli* BL21 (DE3) from a pET-M43 vector in 2YT containing 0.2% glucose. Cultures were grown at 37 °C until an OD_600_ nm of 0.8 was reached and subsequently, protein expression was induced with 0.25 mM IPTG and carried at 18 °C for 16 hours. Cells were harvested and flash frozen in liquid nitrogen and resuspended in lysis buffer (30 mM HEPES/NaOH pH 7.5, 300 mM KCl) for lysis. Target protein was loaded on an amylose column (Dextrin Sepharose™ High Performance, GE Healthcare) and subsequently washed with high-salt buffer (30 mM HEPES/NaOH pH 7.5, 1000 mM KCl) before being re-equilibrated in low-salt buffer (30 mM HEPES/NaOH pH 7.5, 150 mM KCl) and finally eluted in low-salt buffer additionally containing 10 mM D-maltose. The eluted protein was pooled, loaded onto an anion exchange column (HiTrap™ Q FF, Cytiva) and eluted by a linear KCl-gradient (30 mM HEPES/NaOH pH 7.5, 1000 mM KCl). In a final step, eluted protein was polished using a HiLoad™ 16/600 Superdex™ 200 pg (Cytiva) in SEC buffer (20 mM HEPES/NaOH pH 7.5, 150 mM NaCl).

His_6_-TEV-Loc1p constructs (Loc1p^wildtype^, Loc1p^S7C S125C^ and Loc1p^NTR^) and She2p^Dimer^ (She2p^S127E^) were expressed and purified as described previously (Niedner et al. 2013).

### *In vitro* pull-down assays

Proteins were mixed in a total volume of 200 µl pull-down buffer (20 mM HEPES/NaOH pH 7.5, 150 mM NaCl, 2 mM DTT and 2 mM MgCl_2_) at individual concentration of 5 µM. The sample was incubated with 50 µl dextrin sepharose resin (Cytiva Cat. Nr. 28-9355-97) pre-equilibrated in pull-down buffer for 30 min at 4 °C under constant rotation. Beads were washed 4 times with 500 µl pull-down buffer and a finally with 100 µl. Bound proteins were eluted by adding 100 µl of pull-down buffer containing 40 mM D-maltose for 10 minutes at room temperature. Samples for SDS-PAGE analysis were taken from the initially mixed proteins (input, I), from the final wash step (wash, W) and the elution (elu, E). All pull-down experiments were done in triplicates.

### Multi-angle light scattering (MALS)

Proteins were separated on a Superdex™ 200 Increase 10/300 GL column pre-equilibrated in 20 mM HEPES/NaOH pH 7.5, 150 mM NaCl, 2 mM MgCl_2_ and 2 mM DTT with a flow rate of 0.5 ml/min at room temperature. In-line with the size exclusion chromatography proteins were subjected to dynamic light scattering (DLS, DynaPro^®^ NanoStar^®^ II, Wyatt) and multi-angle light scattering (MALS, DAWN^®^ 8, Wyatt) measurements.

### Peptide tiling array

The peptide tiling array with 63 individual peptide spots was ordered at JPT Peptide Technologies. Each peptide has a length of 20 aa and is shifted by 3 aa for each spot. For the detection of She2p-interacting peptides, the membrane was first activated with methanol for 5 minutes at room temperature and blocked with TBS-T containing 1% (w/v) casein (blocking buffer). It was then incubated at room temperature (1) for 16 h at 4 °C with 50 µg/ml She2p diluted in blocking buffer, (2) for 1 h with a primary antibody raised against She2p (1:50 dilution in blocking buffer, SH2 clone 1C3 1-1 rat, CF-MAB Helmholtz Center Munich) and (3) for 1 h with αrat IRDye^®^ 680 RD (1:20.000 dilution in blocking buffer, LI-COR Bioscience). In between the individual incubation steps, the blot was washed 3 times with TBS-T for 5 minutes. To exclude the possibility of detecting unspecific binding of the antibodies to the peptides, a control experiment without She2p was carried out and the membrane was subsequently stripped applying the manufacturers protocol. The membrane was imaged using a LI-COR Odyssey 9120 system.

### Characterization of liquid-liquid phase separated phases

For visualization of LLPS, proteins were mixed in equal ratios at the given concentrations in condensation buffer containing 20 mM HEPES/NaOH pH 7.8, and either 50, 150, or 250 mM NaCl, 2 mM DTT, and 2 mM MgCl_2_. Phases were imaged 5 minutes after mixing the proteins to ensure comparability. Prior to use, object slides and cover slips were treated with 2 % Hellmanex^®^ III for 1 hour, followed by 3 wash steps with H_2_O, a second wash with 1 M NaOH for 30 minutes, and again three wash steps with H_2_O.

To test whether the addition of 5 or 10% of 1,6-hexanediol has an effect on LLPS formation and stability, Loc1p and She2p were mixed equimolar (1.25 or 10 µM) in condensation buffer with 50 mM NaCl containing 5%, 10% or no 1,6-hexanediol (control). Resulting LLPS were imaged as described above.

The droplet area of LLPS at different protein concentrations in 20 mM HEPES/NaOH pH 7.8, 150 mM NaCl, 2 mM DTT and 2 mM MgCl_2_ was quantified using the software FIJI (ImageJ). In a first step, the scale bar in FIJI was defined according to the recorded scale bar on the image. In a second step, the droplets were isolated from the background by setting a threshold and a binary image was created (the same threshold was applied for all analyzed images). The plugin “Analyze particles” was then used to measure the area of the identified droplets.

### Fluorescence labelling and visualization of proteins and nucleic acids in LLPS droplets

Loc1p^S7C S125C^ was labeled using the Cy5 maleimide mono-reactive dye kit (Cytiva Product PA25031) according to the manufacturer’s instructions. After the reaction, the protein was separated from the free dye by applying the centrifuged supernatant to a HiLoad^TM^ 16/600 Superdex^TM^ 75 pg column.

She2p was labeled with Cy3-maleimide (Lumiprobe Product 21080) by first adding 100x excess of TCEP to the protein to reduce potential disulfide bonds for 20 min at RT under nitrogen gas. The Cy3-maleimide was dissolved in DMF to a concentration of 0.5 M and added in 20x excess to the reduced protein. For labeling, the reaction was incubated at 4 °C for 16 h. As for Loc1p^S7C S125C^, the labeled protein was then separated from the free dye by size exclusion chromatography (Superdex^TM^ 75 Increase 10/300 GL column) at a flow rate of 0.5 ml/min. The proteins were flash-frozen in liquid nitrogen and stored at -80 °C. Phase-separated droplets containing labeled protein and/or nucleic acids were imaged at the Core Facility Confocal & Multiphoton Microscopy of Ulm University, using an LSM710 Leica microscope.

Fluorescently labelled E3 ZIP code RNA (5’-Atto390-AUGGAUAACUGAAUCGAAAGACAUUAUCACG) and DNA (5’-Atto390- CCCCCCCTCGAGGTCGACGGTATCGATAAGCTT) were ordered from Biomers. Experiments with RNAs were performed according to published recommendations (Edelmann et al. 2014).

### Chemical crosslinking of She2p and Loc1p

4 μM She2p and 4 μM Loc1p were diluted in activation buffer (0.1 M MES pH 6.0, 300 mM NaCl) and incubated with 8 mM EDC and 20 mM NHS for 15 minutes at room temperature, followed by quenching of EDC by 20 mM 2-mercaptoethanol. The pH was adjusted by adding coupling buffer (0.1 M HEPES pH 8.0, 300 mM NaCl) and the reaction was incubated again for 2 hours at room temperature. The reaction was quenched with 50 mM TRIS pH 8.0 and two negative controls containing only Loc1p or She2p were included. The result was evaluated by SDS-PAGE and western-blot analysis. Subsequently, the proteins were precipitated by adding four-times the reaction volume of 100 % ice-cold acetone followed by thorough mixing. The precipitated sample was stored at -20 °C.

### EDC-crosslink identification by mass spectrometry

The precipitated EDC-crosslinked sample was resuspended in a denaturation buffer (8 M urea, 50 mM ammonium bicarbonate), reduced with 10 mM DTT and alkylated with 55 mM iodoacetamide. The sample was diluted with 50 mM ammonium bicarbonate to reduce urea concentration to 1M and digested with trypsin (Promega, sequencing grade, 1:50 (w/w) enzyme-to-protein ratio) at 37 °C overnight. Obtained peptides was loaded on a C18 micro spincolumn (Harvard Apparatus), washed and eluted twice with 50% acetonitrile (ACN) / 0.1% formic acid (FA) and once with 80% ACN / 0.1% FA. The eluates were pooled and dried in a vacuum concentrator (Eppendorf). The peptides were dissolved in 50 µl of 10 mM ammonium hydroxide and separated by reverse phase HPLC at basic pH using an xBridge C18 3.5µm 1x150mm column (Waters) at a flow rate of 60 µl/min at 24°C. A gradient of buffers A (10 mM ammonium hydroxide, pH 10) and B (80% ACN / 10 mM ammonium hydroxide, pH 10) was used as mobile phase. Peptides were bound to a column pre-equilibrated with 5% buffer B and eluted over 64 min using a multistep gradient of 5-45%B. One-minute fractions (60 µl each) were collected. The fractions corresponding to elution time of 6 to 52 min were pooled in a concatenated manner. The resulting 12 superfractions were vacuum dried and dissolved in 4% ACN / 0.1% TFA for a subsequent uHPLC-ESI-MS/MS analysis. These superfractions were injected into a Dionex UltiMate 3000 uHPLC system (Thermo Scientific) coupled to a Thermo Orbitrap Exploris 480 mass spectrometer and measured in duplicates with a 58 min method. uHPLC was equipped with a PepMAP C18 trap cartridge (0.3 x 5 mm, 5 μm, Thermo Scientific) and a custom 29 cm C18 main column (75 µm inner diameter packed with ReproSil-Pur 120 C18-AQ beads, 3 µm pore size, Dr. Maisch GmbH). Mobile phase was formed using buffers A (0.1% formic acid) and B (80% ACN / 0.08% formic acid). The peptides were separated by applying a linear gradient of 10 to 46%B. MS data were acquired in data-dependent mode using following settings: MS1 resolution, 120000; MS1 scan range, 380-1600 m/z; MS1 normalized AGC target, 300%; MS1 maximum injection time, auto; intensity threshold, 1E4; MS2 resolution, 30000; isolation window, 1.6 Th; normalized collision energy, 28%; MS2 AGC target, 100%; MS2 maximum injection time, 120 ms. Top 30 most abundant precursors with a charge state of 3-8 were selected for MS2 in each cycle. A dynamic exclusion of 30 s was applied.

Protein-protein crosslinks were identified with pLink3.0.17 software (https://pfind.ict.ac.cn/se/plink/) (Chen et al. 2019) by searching Thermo raw files against a database containing sequences of the Loc1p and She2p proteins. Default settings with the EDC-DE preset and false discovery rate (FDR) of 1% were used. The pLink results are shown in **Supplementary Tables 1: and 2**.

### Calculation of the nuclear Loc1p concentration

The number of Loc1p molecules per cell was obtained from the yeast genome database (yeastgenome.org/locus/S000001897, 25.01.2024, 13:50) and the nuclear volume was obtained from (Jorgensen et al., 2007). Using these values and based on the known nuclear localization of Loc1p, the nuclear Loc1p concentration was calculated as follows:

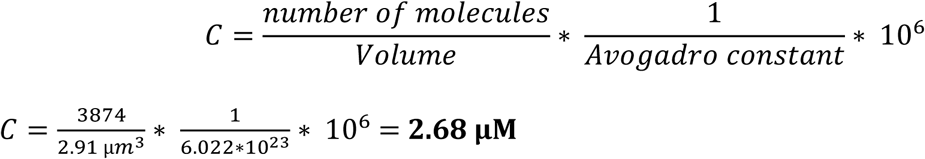

## Acknowledgments and Funding

We like to thank Olexandr Dybkov and Manu Stech Domene for their contribution. H.U. was supported by the DFG (Deutsche Forschungsgemeinschaft) within the Collaborative Research Center SFB860. D.N. was funded by the DFG as part of the Research Unit FOR2333 (NI 1110/5-1 and NI 1110/10-1).

## Supplementary information

**Supplementary Figure 1:**
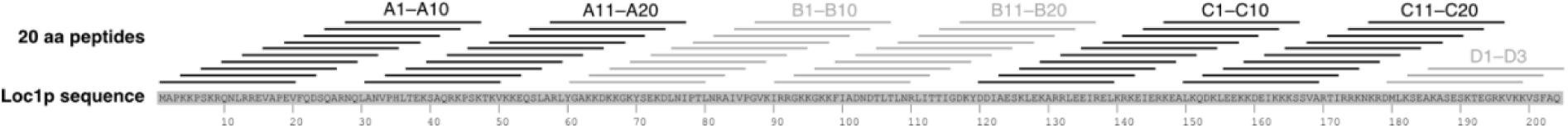
Schematic representation of peptides used for the peptide tiling array aligned with the full Loc1p amino acid sequence.

**Supplementary Figure 2:**
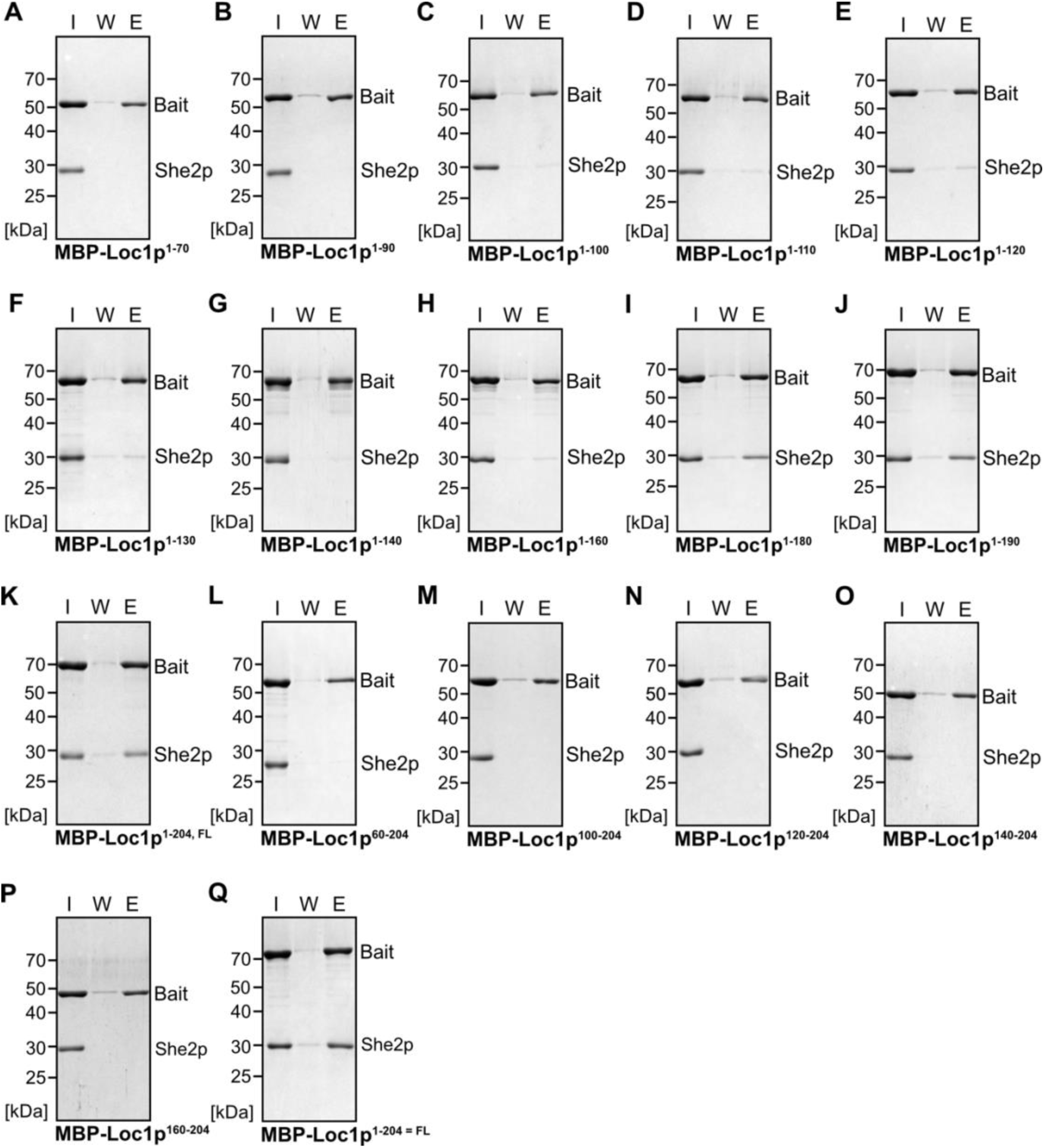
*In vitro* pull-down experiments with all controls. Input- (I), wash- (W), and elution (E) fractions for all pull-down experiments. MBP-fusion proteins were immobilized on amylose beads, and their interaction with She2p at 5 µM input concentration was tested. All pull-downs were done in triplicates.

**Supplementary Figure 3:**
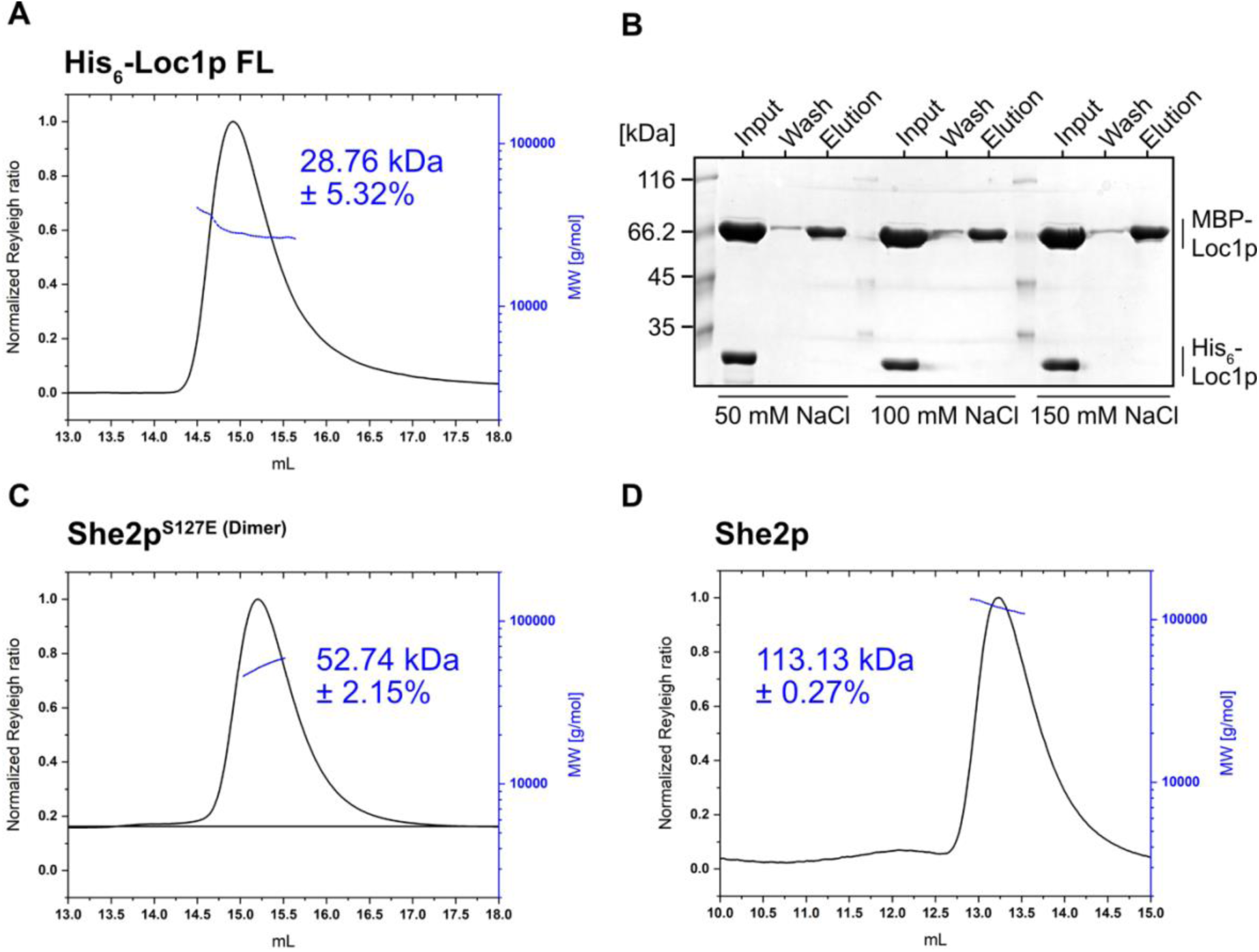
Analysis of the oligomeric states of His_6_-Loc1p and She2p variants. Multi-angle light scattering (MALS) measurements at 115 µM **(A)** and pull-down experiments with 5 µM Loc1p concentrations **(B)** show that Loc1p is monomeric in solution. For monomeric His_6_-Loc1p a theoretical molar mass of 26.0 kDa was calculated, which agreed well with the experimentally determined value of 28.8 kDa. MALS measurements with She2p^S127E^ **(C)** and She2p **(D)** show that their oligomeric states differ. The determined molar mass of 113.1 kDa for She2p closely matches the theoretical value of 114.8 kDa for a tetramer (28.8 kDa per monomer). In contrast, the phosphomimetic mutant She2p^S127E^ forms a dimer in solution. Efforts to determine the molar mass or stoichiometry of the complex did not yield in interpretable data (not shown).

**Supplementary Figure 4:**
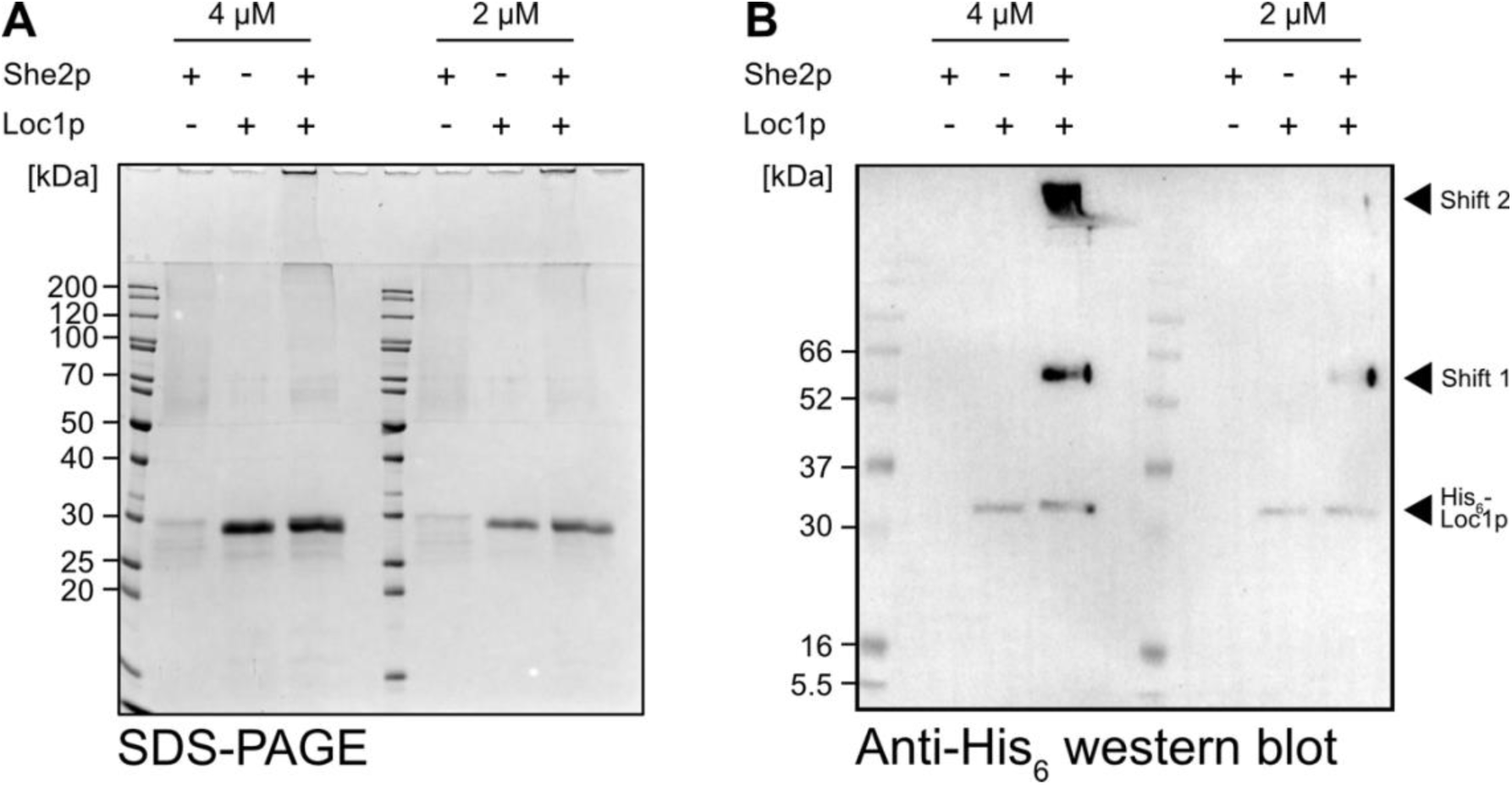
Cross-linking with She2p and His_6_-Loc1p. One-step EDC/NHS chemical cross-linking experiments with recombinant She2p and His_6_-Loc1p, which were performed to scout for suitable protein concentrations and experimental conditions. She2p and Loc1p were crosslinked individually as control, and together at equimolar concentrations (either 4 μM or 2 μM). Bands and band shifts were analyzed by SDS-PAGE **(A)** and a western blot **(B)** against the His_6_-tag of His_6_-Loc1p. Arrows indicate band shifts containing His_6_-Loc1p as well as His_6_-Loc1p-containing cross-linked assemblies or complexes (presumably with She2p).

**Supplementary Table 1:** She2p Crosslinks to Loc1p.

| **Position (She2p)** | **Amino acid** | **Count of peptides** |
| --- | --- | --- |
| 3 | Lys | 15 |
| 15 | Glu | 6 |
| 18 | Glu | 40 |
| 67 | Asp | 10 |
| 76 | Glu | 30 |
| 80 | Glu | 5 |
| 82 | Lys | 22 |
| 84 | Glu | 88 |
| 86 | Asp | 23 |
| 88 | Glu | 30 |
| 93 | Asp | 10 |
| 132 | Glu | 127 |
| 133 | Asp | 6 |
| 139 | Glu | 23 |
| 170 | Asp | 8 |
| 227 | Asp | 61 |
| 228 | Glu | 640 |
| 229 | Glu | 153 |
| 231 | Asp | 55 |
| 236 | Lys | 11 |
| 240 | Lys | 14 |
| 243 | Lys | 13 |

**Supplementary Table 2:** Loc1p crosslinks to She2p.

| **Position (Loc1p)** | **Amino acid** | **Count of peptides** |
| --- | --- | --- |
| 15 | Glu | 22 |
| 19 | Glu | 12 |
| 38 | Glu | 11 |
| 39 | Lys | 73 |
| 44 | Lys | 4 |
| 47 | Lys | 16 |
| 51 | Lys | 65 |
| 52 | Lys | 116 |
| 53 | Glu | 4 |
| 63 | Lys | 40 |
| 69 | Lys | 82 |
| 73 | Lys | 82 |
| 74 | Asp | 9 |
| 89 | Lys | 21 |
| 98 | Lys | 113 |
| 102 | Asp | 4 |
| 117 | Asp | 3 |
| 118 | Lys | 76 |
| 121 | Asp | 6 |
| 126 | Lys | 69 |
| 129 | Lys | 13 |
| 142 | Lys | 9 |
| 147 | Lys | 36 |
| 151 | Lys | 81 |
| 154 | Lys | 25 |
| 158 | Lys | 18 |
| 159 | Lys | 4 |
| 163 | Lys | 22 |
| 164 | Lys | 12 |
| 165 | Lys | 100 |
| 182 | Lys | 84 |
| 186 | Lys | 52 |
| 191 | Lys | 47 |
| 199 | Lys | 55 |

